# NOC1 is a Direct MYC Target, and Its Protein Interactome Dissects Its Activity in Controlling Nucleolar Function

**DOI:** 10.1101/2023.09.12.557489

**Authors:** Valeria Manara, Marco Radoani, Romina Belli, Daniele Peroni, Francesca Destefanis, Luca Angheben, Gabriele Tome, Toma Tebaldi, Paola Bellosta

**Affiliations:** Department of Computational, Cellular, Integrative Biology CIBIO, University of Trento 38123, TN, Italy; Evolutionary Biology CSIC Universitat Pompeu Fabra, 08003 Barcelona, Spain; Department of Internal Medicine, Yale School of Medicine, 06520, New Haven, CT, USA; Department of Medicine, NYU Langone Medical Center, 10016 New York, USA

**Keywords:** NOC1, MYC, E-box, nucleolus, Mass Spectrometry, Interactome

## Abstract

The nucleolus is a subnuclear compartment critical in ribosome biogenesis and cellular stress responses. These mechanisms are governed by a complex interplay of proteins, including NOC1, a member of the NOC family of nucleolar proteins responsible for controlling rRNA processing and ribosomal maturation. This study reveals a novel relationship between NOC1 and MYC transcription factor, known for its crucial role in controlling ribosomal biogenesis, cell growth, and proliferation. Here, we demonstrate that NOC1 functions as a direct target of MYC, as it is transcriptionally induced through a functional MYC-binding E-box sequence in the NOC1 promoter region. Furthermore, protein interactome analysis reveals that NOC1-complex includes the nucleolar proteins NOC2 and NOC3 and other nucleolar components such as Nucleostemin1 Ns1 transporters of ribosomal subunits and components involved in rRNA processing and maturation. In response to MYC, NOC1 expression and localization within the nucleolus significantly increase, suggesting a direct functional link between MYC activity and NOC1 function. Notably, NOC1 over-expression leads to the formation of large nuclear granules and enlarged nucleoli, which co-localize with nucleolar fibrillarin and Ns1. Additionally, we demonstrate that NOC1 expression is necessary for Ns1 nucleolar localization, suggesting a role for NOC1 in maintaining nucleolar structure. Finally, the co-expression of NOC1 and MYC enhances the formation of abnormal structures formed by NOC1 within the nucleolus, outlining another aspect of NOC1 and MYC cooperation in nucleolar dynamics.

This study also reveals an enrichment with NOC1 with few proteins involved in RNA processing, modification, and splicing. Moreover, proteins such as Ythdc1, Flacc, and splenito are known to mediate N6-methyladenosine (m6A) methylation of mRNAs in nuclear export, revealing NOC1’s potential involvement in coordinating RNA splicing and nuclear mRNA export.

In summary, we uncovered novel roles for NOC1 in nucleolar homeostasis and established its direct connection with MYC in the network governing nucleolar structure and function. These findings also highlight NOC1’s interaction with proteins relevant to specific RNA functions, suggesting a broader role in addition to its control of nucleolar homeostasis and providing new insight that can be further investigated.

## Introduction

MYC is a transcription factor crucial in the regulation of factors controlling ribosomal biogenesis and protein synthesis, which occurs primarily through its ability to regulate the transcription of genes required for ribosome assembly and function (van RiggelenYetil and Felsher, 2010;Campbell and White, 2014;DestefanisManara and Bellosta, 2020). MYC promotes the transcription of its target genes, such as ribosomal proteins and co-factors, by binding to specific DNA sequences known as E-boxes (5’-CACGTG-3’) within their promoter region (Fernandez et al., 2003;Orian et al., 2003;Hulf et al., 2005). MYC also promotes the transcription of ribosomal RNA (rRNA) genes, which are transcribed by RNA polymerase I to generate the precursor rRNA transcripts. Since ribosomes are central to protein synthesis and cell growth, MYC’s role in promoting ribosomal biogenesis largely contributes to protein synthesis necessary for cell growth and proliferation, a function that is conserved both in flies and vertebrates (Schlosser et al., 2003;Arabi et al., 2005;Grandori et al., 2005;Grewal et al., 2005;van RiggelenYetil and Felsher, 2010;DestefanisManara and Bellosta, 2020).

NOC1 is a nucleolar protein that, together with NOC2 and NOC3, plays a critical role in the maturation of rRNA and the transport of the pre-ribosomal subunits (Sailer et al., 2022;Dorner et al., 2023). NOC1 in yeast works as a heterodimer with NOC2 during the initial transport and maturation of the ribosomal RNA (rRNA), a process that is completed by NOC2/NOC3 heterodimers (Milkereit et al., 2001). Furthermore, studies on the distribution of affinity-tagged NOC1 and, more recently, proteomics and crosslinking coupled to mass spectrometry confirmed the presence of NOC1 in the early pre-60S complex (Sailer et al., 2022;Dorner et al., 2023), while cryo-EM studies showed its role in the formation of heterodimers with NOC2, essential for the quality-control checkpoint of the maturation of the large ribosome subunit (Sanghai et al., 2023).

We recently characterized NOC1 function in flies and showed its role in controlling polysome abundance, rRNA maturation, protein synthesis, and cell survival (Destefanis et al., 2022). Furthermore, lowering NOC1 levels in different contexts, such as whole animals or specific organs, results in various developmental and functional impairments (Destefanis et al., 2022). Our initial transcriptomic analysis revealed NOC1 as a potential direct target of MYC (Hulf et al., 2005); thus, we further analyzed this critical function in the context of ribosomal biosynthesis directed by MYC.

Here, we show that NOC1 is a direct transcriptional target of MYC, and its activation is mediated by a functional E-box sequence located in the promoter region of the *NOC1* gene. We then used HA-NOC1 as bait to perform Mass Spectrometry (MS) analysis to determine the NOC1 interactome to characterize NOC1 function and connect its activity with biological processes, mainly focusing on components that control nucleolar homeostasis.

Bioinformatic analysis using the STRING database identified clusters of NOC1 protein interactors, and the most significant was on ribosome biogenesis. These data showed a significative enrichment of NOC2 and NOC3 (*p*<0.05) strongly aligning with data published previously in yeast (Milkereit et al., 2001;Hierlmeier et al., 2013), and a significant cluster of nucleolar proteins, such as fibrillarin (fib) and nucleostemin 1 (Ns1), and others, like Novel nucleolar proteins (Non1 and Non3) and mushroom body miniature (mbm) involved in the 60S subunit biogenesis. Moreover, we found an enrichment of nucleolar and nuclear proteins, like Nnp (Hulf et al., 2005), and peter pan (ppan) (MigeonGarfinkel and Edgar, 1999;Zielke et al., 2022), involved in pre-rRNAs and controlling RNA maturation, and modulo (mod) (Perrin et al., 2003), that were previously identified as direct targets of MYC, emphasizing the relation between NOC1 and MYC.

In addition, these studies also identified enrichment of the nuclear m^6^A ‘reader’ YTH domain RNA Binding Protein C1 (Ythdc1) (Roundtree et al., 2017), Flacc (Fl(2)d-associated protein), and spenito (nito)(Knuckles et al., 2018). Remarkably, these proteins are part of the complex that mediates the N6-Methyladenosine methylation of mRNAs for their nuclear export (Knuckles et al., 2018;Shi et al., 2021). We could outline a novel function for the MYC-NOC1 axis in regulating mRNA m^6^A modification and transport.

Finally, the observation that NOC1 controls the nucleolar localization of Ns1, together with those indicating that MYC enhances NOC1-induced large granular structures in the nucleus, further sustains the functional relationship between MYC and NOC1 in maintaining nucleolar homeostasis.

In summary, these findings will provide significant insights into the role of NOC1 and its interactome that may contribute to the control of nucleolar functions, supporting the crucial role of MYC in regulating growth, proliferation, and protein synthesis.

## Materials and Methods

### Fly stocks and husbandry

Fly cultures and crosses were raised at 25 °C on a standard medium containing 9 g/L agar (ZN5 B and V), 75 g/L corn flour, 60 g/L white sugar, 30 g/L brewers’ yeast (Fisher Scientific), 50 g/L fresh yeast and 50 mL/L molasses (Naturitas), along with nipagin and propionic acid (Fisher). The lines used were obtained by: *UAS-HA-MYC (Bellosta et al., 2005); NOC1-GFP (B51967) UAS-NOC1-HA* (Flyorf-CH) *NOC1-RNAi* (B25992). *UAS-Ns1-GFP* is a gift from Patrick J. Di Mario University of Louisiana, LA). *hsp70-Gal4* gift from Florenci Serras (University of Barcelona, Spain).

### Cloning NOC1 E-box and Molecular biology

Site-directed mutagenesis (SDM) was carried out using the following primers for the mutant E-box 5’ TTC GGC ACG AGT TTG AAT AGA ATT CCG AGT TGT TTC TAA CGC CG; 5’ CGG CGT TAG AAA CAA CTC GGA ATT CTA TTC AAA CTC GTG CCG AA; following instructions from the SDM kit (Promega). Promoter elements used in luciferase reporter expression analyses were cloned into the pGL3-basic vector (Promega).

### Cell culture and luciferase assays

S2 *Drosophila* cells were propagated in Schneider’s Drosophila medium (Gibco), supplemented with 10% fetal bovine serum, at 24°C. S2 cell transfections were carried out using Cellfectin (Invitrogen). *NOC1* reporter constructs were added at 1 µg per 10^6^ cells; tubulin-Renilla luciferase control DNA were co-transfected at 0.1 µg per 10^6^ cells and incubated with a transfection mix for 12 h. Cells were harvested 24 or 60 h posttransfection. Relative gene expression was determined using the Dual-Luciferase Reporter assay system (Promega) on a luminometer.

### RNA extraction and quantitative RT-PCR analysis

Total RNA was extracted from 8 whole larvae using the QIAGEN RNeasy Mini Kit (Qiagen) according to the manufacturer’s instructions. Extracted RNAs were quantified using an ultraviolet (UV) spectrophotometer, and RNA integrity was confirmed with ethidium bromide staining. 1 μg total RNA from each genotype was reverse transcribed into cDNA using SuperScript IV MILO Master Mix (Invitrogen). The obtained cDNA was used as the template for quantitative real-time PCR (qRT-PCR) using qPCR Mastermix (Promega). mRNAs expression levels were normalized to *actin-5C mRNA* used as the internal control. The relative level for each gene was calculated using the 2-DDCt method (Hulf et al., 2005) and reported as arbitrary units. Three independent experiments were performed and cDNAs were used in triplicate. The following primers were used for qRT-PCR: *Actin5c:* 5’CAGATCATGTTCGAGACCTTCAAC; 5’ACGACCGGAGGCGTACAG (Parisi et al., 2013). *Fibrillarin:* 5’ACGACAGTCTCGCATGTGTC; 5’ATGCGGTACTTGTGTGGATG (this work). *MYC*: 5’CATAACGTCGACTTGCGTG; 5’GAAGCTCCCTGCTGATTTGC (Parisi et al., 2013). *NOC1:* 5’CTATACGCTCCACCGCACAT; 5’GTCGCTACCGAACTTGTCCA (Destefanis et al., 2022).

### Protein extractions and western blotting

Larvae 5 for each genotype were lysed in 200 μl of lysis buffer (50 mM Hepes/pH 7.4, 250 mM NaCl, 1 mM (EDTA), 1.5% Triton X-100 containing a cocktail of phosphatases inhibitors (PhosSTOP 04906837001, Merck Life Science) and proteases inhibitors (Roche, cOmplete Merck Life Science). Samples were sonicated three times for 10 seconds using a Branson Ultrasonic Sonifier 250 (Branson, Darbury, CA, USA) equipped with a microtip set at 25% power. Tissue and cell debris were removed by centrifugation at 10,000× *g* for 30 min at 4 °C. Proteins in the crude extract were quantified by a bicinchoninic acid (BCA) Protein assay Reagent Kit (Pierce), following the manufacturer’s instructions with bovine serum albumin as the standard protein. For SDS-PAGE, samples were incubated for 8 min at 100 °C in standard reducing 1× loading buffer; 40 µg of total protein were run on an SDS-polyacrylamide gel and transferred onto nitrocellulose membranes (GE-Healthcare, Fisher Scientific Italia) After blocking in 5% (*w*/*v*) non-fat milk in tris-buffered saline (TBS)-0.05% Tween (TBS-T), membranes were incubated overnight with primary antibodies: rat monoclonal anti-HA (1:1000, ROCHE), or Actin5c (1:200, #JL20) from Developmental Studies Hybridoma Bank (DSHB), University of Iowa, IA, USA. Appropriate secondary antibody was incubated for 2 hours at room temperature, followed by washing. The signal was revealed with ChemiDoc Touch Imaging System (Bio-Rad Lab).

### Immunoprecipitation

*Hsp70 (hs)-Gal4> NOC1* larvae or control *hs-Gal4> w^1118^* were heat-shocked at 37 °C for one hour and left to recover for two hours at room temperature. 20 larvae from each genotype were washed in PBS and lysed with 750 µl of immunoprecipitation buffer (100mM HEPES, 100mM NaCl, 0.5% Triton, 10mM MgCl) containing proteases and phosphatases inhibitors. Protein lysates were incubated for 20 minutes in ice and centrifuged at 13.000 rpm for 30 minutes a 4 °C. 500 µl of lysates were incubated with 50 µl of Sepharose-beads-Protein-G (Invitrogen) previously incubated with 4 µl anti-HA antibodies. Incubation was performed for 2 hours at room temperature, and beads were washed extensively with ice cold lysing buffer. After centrifugation, bound proteins were eluted with 100 µl of SDS-loading buffer LDS Sample Buffer (Thermo Fisher Scientific) containing 5% Bolt Sample reducing agent (Thermo Fisher Scientific) at 80°C for 5 min. 20 µl of the sample was run on a Western blot and 80 µl were used for the MS analysis. Experiments were repeated twice.

### Mass Spectrometry and Proteomic interaction partners analysis

Immunoprecipitated samples were loaded on 10% SDS-PAGE and run for about 1 cm. Gels were then stained with Coomassie and the entire stained area was excised as one sample. Excised gel bands were cut into small plugs (∼1mm3), rinsed with 50 mM ammonium bicarbonate and acetonitrile (ACN) solution, and vacuum dried. Dried gel pieces were then reduced using 10mM DTT (56°C for 30 min) and alkylated using 55mM iodoacetamide (room temperature for 30 min, in the dark). After sequential washing with 50 mM NH4HCO3 and ACN, gel pieces were dried and rehydrated with 12.5 ng/mL trypsin (Promega, Madison, WI) solution in 25 mM ammonium bicarbonate on ice for 30 min. The digestion was continued at 37°C overnight. The tryptic peptides were sequentially extracted from the gels with 30% ACN/3% TFA and 100% ACN. All of the supernatants were combined and dried in a SpeedVac. The tryptic peptides were resuspended in 0.1% TFA, desalted on C18 stage tips, and resuspended in 20 μl of 0.1% formic acid buffer.

For LC-MS/MS analysis, the peptides were separated on an Easy-nLC 1200 UHPLC system (Thermo Fisher Scientific) using an 85-minute gradient on a 25 cm long column (75 µm inner diameter) filled in-house with C18-AQ ReproSil-Pur material (3 µm particle size, Dr. Maisch, GmbH). The gradient was set as follows: from 5% to 25% in 52 minutes, from 25% to 40% in 8 minutes, and from 40% to 98% in 10 minutes, with a flow rate of 400 nL/min. The buffers were 0.1% formic acid in water (A) and 0.1% formic acid in acetonitrile (B). The peptides were analyzed with an Orbitrap Fusion Tribrid mass spectrometer (Thermo Fisher Scientific, San Jose, CA, USA) in data-dependent mode. Full scans were performed in the Orbitrap mass analyzer at a resolving power of 120,000 FWHM (at 200 m/z) in the mass range of 350-1100 m/z, with a target value of 1×106 ions and a maximum injection time of 50 ms. Each full scan was followed by a series of MS/MS scans (collision-induced dissociation) over a cycle time of 3 seconds, with a maximum injection time of 150 ms (ion trap) and a target of 5×103 ions. The ion source voltage was set at +2100V and the ion transfer tube was warmed up to 275°C. Data was acquired using Xcalibur 4.3 and Tune 3.3 software (Thermo Fisher Scientific). QCloud was used for all acquisitions to control instrumental performance during the project, using quality control standards (Chiva et al., 2018).

For data and computational analysis, the raw files were searched in Proteome Discoverer version 2.2 software (Thermo Fisher Scientific). Peptide searches were performed using the UniProt Drosophila melanogaster (fruit-fly) database digested in silico (downloaded in July 2022) and a database containing common contaminants. Trypsin was chosen as the enzyme with 5 missed cleavages. The static modification of carbamidomethylation (C) was incorporated in the search, with variable modifications of oxidation (M) and acetylation (protein N-term). The MASCOT search engine (v.2.2 Matrix Science) was used to identify the proteins, using a precursor mass tolerance of 10 ppm and a product mass tolerance of 0.6 Da. False discovery rate was filtered for <0.01 at PSM, at peptide and protein levels. Results were filtered to exclude potential contaminants and proteins with less than two peptides.

MS downstream analysis was performed using the ProTN proteomics pipeline (www.github.com/TebaldiLab/ProTN and www.rdds.it/ProTN) (manuscript in preparation). Peptide intensities were log2 transformed, normalized (median normalization), and summarized into proteins (median sweeping) with functions in the DEqMS Bioconductor package (Zhu et al., 2020). Imputation of the missing intensities was executed by PhosR package (Kim et al., 2021). Differential analysis was performed with the DEqMS package, proteins with absolute log2 FC > 0.75 and p-value < 0.05 were considered significant.

Protein-protein interaction network was constructed using STRING interaction database, version 12.0 (https://string-db.org/) (von Mering et al., 2003). Medium confidence interactions (score>0.4) were accepted as determined by the STRING database. The PPI network was grouped into relevant protein clusters using the Markov Cluster Algorithm (inflation parameter, 3) clustering option provided by STRING.

### Immunostaining

Dissected tissues were fixed in 4% paraformaldehyde (PFA) (Electron Microscopy Science) in PBS for 30 minutes at room temperature. After permeabilization with 0.3% Triton/PBS, tissues were washed in Tween 0.04% in PBS, saturated with 1% BSA in PBS, and incubated overnight with anti-fibrillarin antibodies (1:100), anti-HA (1:100, ROCHE), anti-GFP (1:200, ThermoFisher A11122) and anti-MYC affinity-purified antibodies (1:1000) (Galletti et al., 2009;Destefanis et al., 2022). Relative secondary antibodies conjugated with Alexa555 were used 1:2000 (Invitrogen). After washing with PBST, samples were mounted on slides using Vectashield (Vector Laboratories) and fluorescence images were acquired using a Leica-CS-SP8 confocal microscope.

## Results

### NOC1 contains a functional E-box sequence in its promoter region and is transcriptionally induced by MYC

Our initial observation on the transcriptomic analysis of potential MYC target genes identified NOC1 as a predicted nucleolar gene that contains in its 5’ promoter region the E-box sequence CACGTG typically within the first 100 bp from the initial translation initiation codon ATG (Figure 1A), and thus considered a *bona-fide* MYC binding region (Hulf et al., 2005). By qRT-PCR, we show that constitutive expression of MYC in whole *Drosophila* larvae (Figure 1B) using the *actin* promoter resulted in NOC1 transcriptional activation and also in the upregulation of *fibrillarin-mRNA* (Figure 1C), a known MYC target that contains functional E-boxes in its promoter region conserved both in flies and vertebrates (Orian et al., 2003;Hulf et al., 2005;Koh et al., 2011).

**Figure 1.**
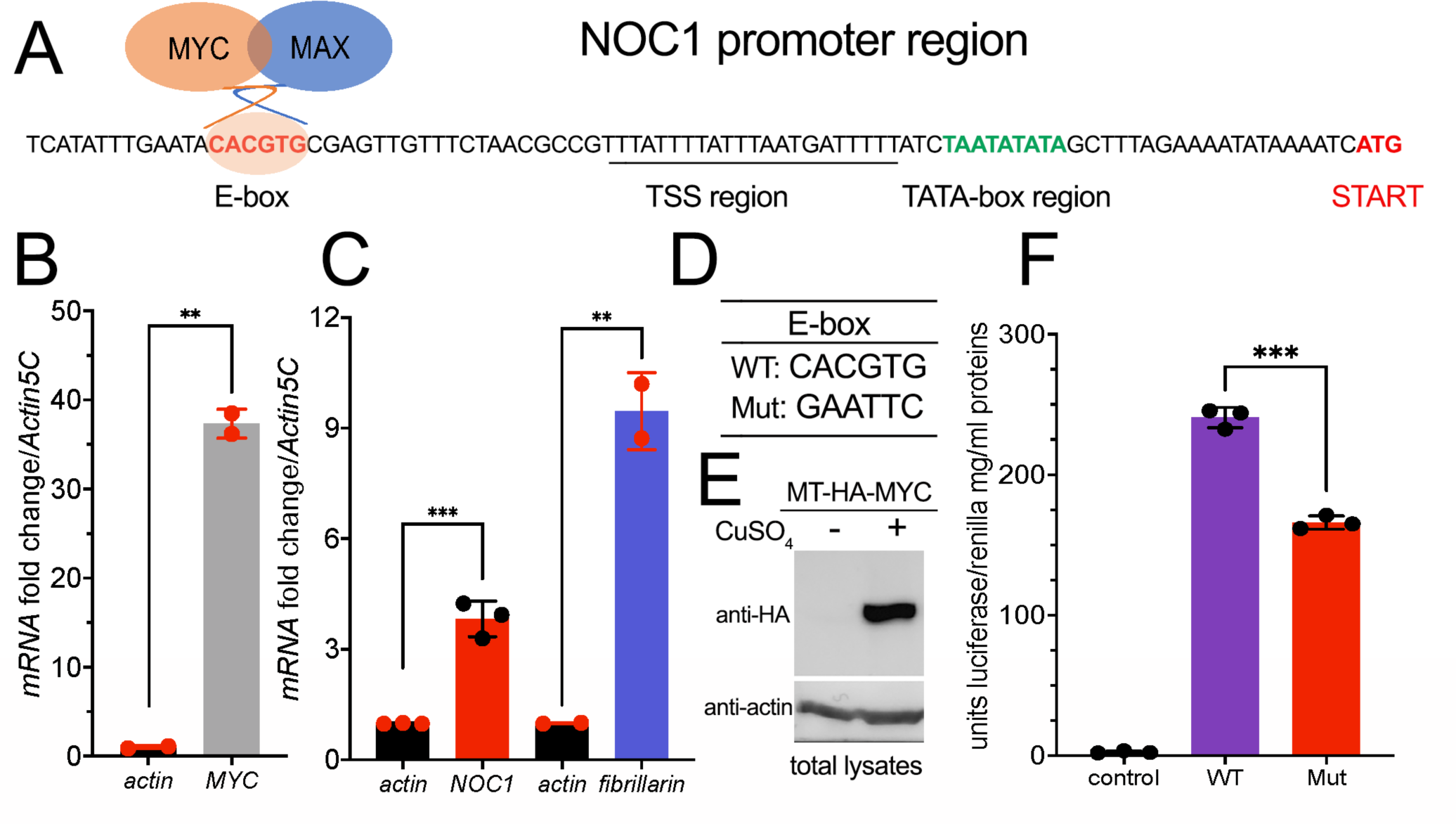
NOC1 contains in its promoter a functional MYC E-box sequence. (A) DNA promoter region of the *NOC1* gene showing the position of the E-box, the putative Transcription Start Sequence (TSS) with the TATA box, and the Initiation of Transcription point (START). (B) qRT-PCR from third instar whole larvae tissues showing the upregulation of *MYC-mRNA* (B) and of *NOC1* and *fibrillarin-mRNAs* (C) upon MYC induction. The expression of *UAS-MYC* was induced using the *actin-Gal4* promoter. (D) DNA sequences of WT and Mutant E-boxes. (E) Western blot from S2-MT-HA-MYC cells showing the expression level of the metallothionein HA-MYC upon induction for five hours using CuSO_4._ Actin was used as a control for loading. (F) Units of relative luciferase activity in lysates of S2-MT-HA-MYC cells treated for five hours with CuSO_4_ and transfected with Renilla plasmid alone (control), of with NOC1 promoter region containing WT (WT) or Mutant (Mut) E-box.

The 5’ promoter region of NOC1 contains a putative TATA box sequence at about - 26 bp from the transcription start, a sequence identified as the Transcription Start Site (TSS), and the CACGTG sequence (E-box) at -82 bp from the ATG transcription start (Figure 1A). To investigate whether the CACGTG sequence responds to MYC activation, we cloned the 5’ promoter region of *NOC1*, containing the wildtype CACGTG sequence or the scramble sequence GAATTC (Figure 1D), upstream of a plasmid expressing the Firefly luciferase ORF. The reporter plasmids were co-transfected into *Drosophila* S2-MT-MYC cells with a plasmid expressing the Renilla luciferase. MYC expression was induced by adding CuSO4 to the medium (Figure 1E). Firefly luciferase activity was measured in the cell lysates after five hours of induction and normalized to the co-transfected Renilla luciferase expressed under the control of the constitutive tubulin promoter (Figure 1F). As shown upon MYC expression, cells expressing the *NOC1* promoter region with the mutated E-box have significantly reduced luciferase activity compared to that from cells expressing the wild-type *NOC1* promoter, indicating that the sequence CACGTG in the *NOC1* promoter functions as an enhancer of MYC activity.

### Interactome analysis of NOC1 associates its expression with NOC2 and NOC3 proteins and other components of the nucleolus

To investigate how NOC1 might regulate nucleolus functions, we explored its binding partners by analyzing the total interactome through immunoprecipitation and tandem mass spectrometry analysis (Figure 2A). Third-instar larvae expressing *UAS-HA-NOC1* under the *actin-Gal4* were used first to test a few conditions to efficiently extract NOC1 protein from the cells (Figure 2B). As shown in the left panel, NOC1 is efficiently expressed in lysates from third-instar larvae as a 120 KDa protein detected by the anti-HA antibodies. We first tested three conditions for lysing the tissues to avoid high detergent and salt concentrations according to previous protocols for immunoprecipitation in whole larvae (Bellosta et al., 2005). Since all three conditions used gave an excellent recovery of NOC1 in the tissue lysates, we decided to pursue our experiments using a lysing buffer that contained 0.5% Triton and 200 mM NaCl (Figure 2; middle panel). Since we found NOC1 transcriptionally upregulated as early as 3 hours upon MYC expression (Hulf et al., 2005) and (Figure 1C), we decided to use the inducible promoter *hsp70 (heat-shock)-Gal4* to ubiquitously express NOC1 to perform our analysis at a similar time point. *Hs-Gal4; UAS-HA-NOC1* larvae and control (*hs-Gal4; w^1118^*) were heat-shocked for one hour and 37 °C. After two hours of recovery at room temperature, larvae were lysed to pursue the immunoprecipitation (IP) using anti-HA antibodies. Immunoblotting analysis showed enrichment of HA-NOC1 bands in the expected samples (Figure 2; left panel). While a weak band of 120 KDa is also visible in the control sample, the lower molecular weight bands characteristic of the NOC1 pattern are not present (Destefanis et al., 2022), confirming the specificity of the experiment.

**Figure 2:**
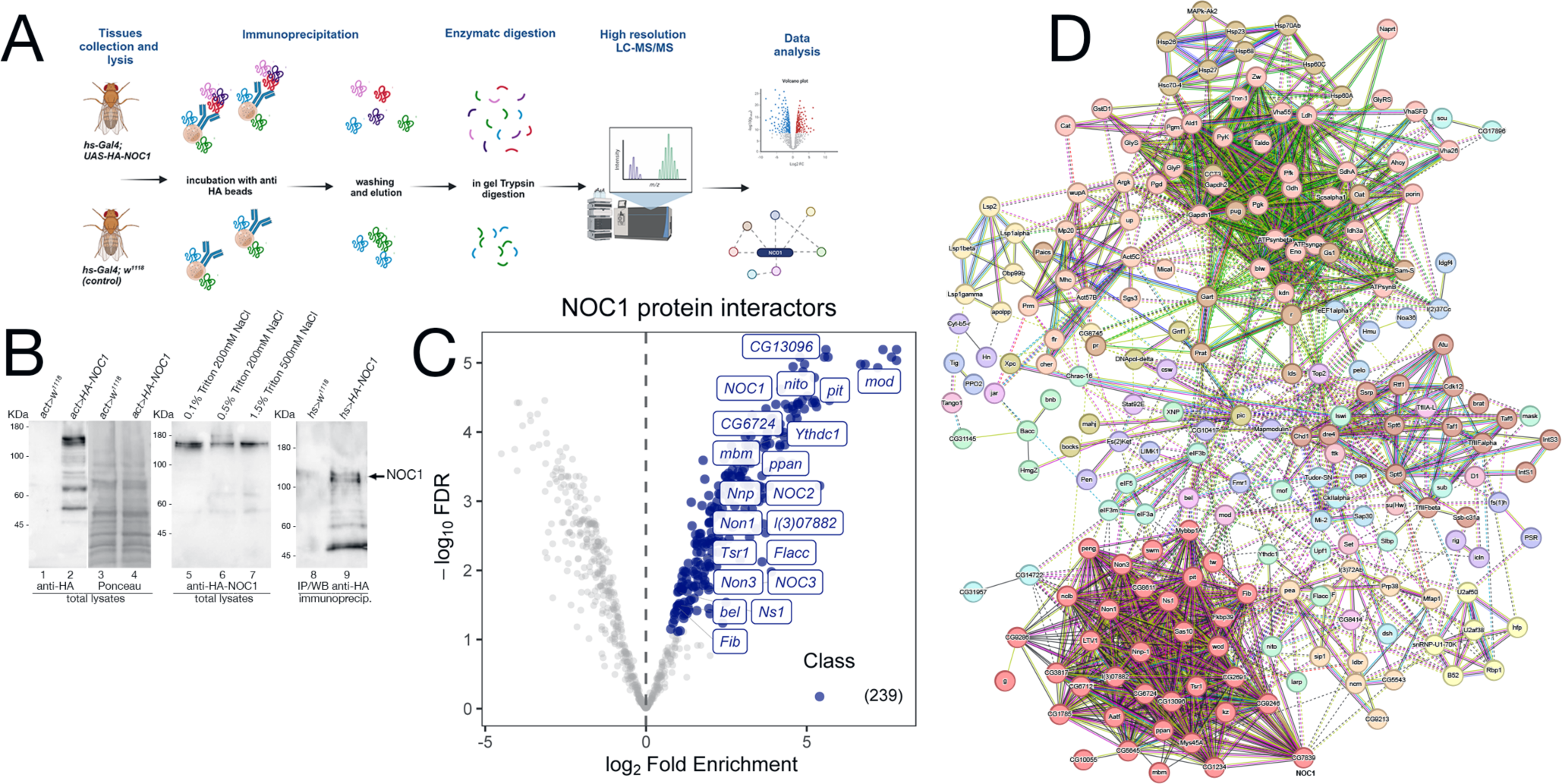
NOC1 is associated with components of the nucleolus. (A) Schematic representation of the workflow used to identify NOC1 interacting proteins. *hs-Gal4; UAS-HA-NOC1* larvae and control (*hs-Gal4; w^1118^*) were lysed and subjected to immunoprecipitation using anti-HA conjugated beads. NOC1 immunoprecipitated proteins were eluted with Laemmli sample buffer and processed by *in-gel trypsin* digestion before MS/MS analysis. The figure is created with BioRender.com. (B) Western blot showing the expression of *UAS-HA-NOC1* in lysates from third instar larvae and the enrichment in the IPs using the *actin>Gal4* promoter. The left panel shows a band of about 120KDa recognized by anti-HA antibodies and present only in the total lysates of control larvae (lane 1) or expressing HA-NOC1 (lane 2). In lanes 3 and 4 is shown the Ponceau staining relative to lanes 1 and 2. In the middle panel is shown the expression level of HA-NOC1 upon immunoblot with anti-HA antibodies from larvae lysate with buffer containing different concentrations of detergent and salt (lane 5-7). In the right panel is shown the immunoblot from the eluted material from the Seph-Prot-G conjugated with anti-HA antibody upon immunoprecipitation from lysates of larvae expressing HA-NOC1 using the *hsp70-heat-shock (hs)* inducible promoter after one hour of heat-shock and two hours of recovery. Lane 8 shows the immunoblot from lysates of control larvae *hs-w^1118,^* while lane 9 shows the eluted from the immunoprecipitation from animals expressing *hs-HA-NOC1*; this represents 1/5 of the material used from the MS analysis. (C) Volcano plot highlighting all proteins enriched. The mean log2 ratio of hs-HA-NOC1 IPs versus control hs-w^1118^ IPs are plotted versus the corresponding *p*-values. 239 proteins significantly enriched (blue dots) with a *p*-value below 0.05 and log2-FC >1.5 thresholds were treated as putative NOC1 binding partners. The most representative interactors found for this analysis are indicated in the plot. (D) Schematic view of protein-protein interactions among NOC1 targets according to the STRING database (v.12). STRING protein-protein interaction analysis indicates the most prominent clusters with a medium confidence score of 0.4. Each node represents a protein, and each edge represents an interaction.

To discover NOC1 protein partners, we used affinity purification coupled with label-free mass spectrometry (AP-MS). Specifically, we performed the co-immunoprecipitation of the tagged-NOC1 protein in *hs-Gal4; UAS-HA-NOC1* lysates, and the control tissues *hs-Gal4; w^1118^*, respectively. Immunoprecipitates (IPs) were then analyzed by LC-MS/MS using an Easy-nLC 1200 UHPLC system coupled to an Orbitrap Fusion™ mass spectrometer. For protein identification and quantification, acquired raw data were imported into the Proteome Discoverer 2.2 (PD) platform and searched with MASCOT (v2.6 Matrix Science, London, U.K.) against the UniProtKB Drosophila melanogaster database. The quantitative output of PD was then further processed using the ProTN pipeline, enabling comprehensive quality control, statistical analysis, and interpretation of proteomic datasets. We identified a total of 239 proteins that were significantly (p<0.05) enriched in HA-NOC1 immunoprecipitated (IPs) relative to control, representing putative NOC1 binding partners (Supplementary List1). Results are illustrated by the volcano plot displaying the proteins significantly enriched in NOC1-IPs in light blue, with a fold change (FC) > 1.5 and p-value < 0.05. To better characterize the NOC1 interactome, the list of putative interacting proteins was processed by STRING protein-protein interaction analysis, and clusters were identified in the resulting network using the Markov Clustering Algorithm (MCL) (Figure 2C). This analysis outlined a few interesting clusters of NOC1 interactors (Figure 2D). The most relevant is Cluster1, which includes NOC2 and NOC3 (Supplementary Table1, and Volcano plot Figure 2C). The same cluster also includes nucleolar proteins such as Fibrillarin (Fib), an rRNA O-methyltransferase, and l(3)07882 required for the processing of the pre-rRNAs, Novel nucleolar proteins (Non1 and Non3) involved in the biogenesis of the 60S subunits and needed for the assembling of the mitotic spindle, like Nucleostemin 1 (Ns1), required for the release of the 60S ribosomal subunit, mushroom body miniature (mbm) involved in ribosome biogenesis. Others not nucleolar proteins, like the CG13096, a homolog of human Ribosomal L1 domain-containing protein (RSLD1), the CG6724, a putative homolog of WRD12 required for the maturation of rRNAs and the formation of the large ribosomal subunit, Nnp, and Tsr1 described for the processing of pre-rRNAs and the control of RNA maturation. Notably, we also found in the interactome the DEAD-box RNA helicases pitchoune (pit) (Zaffran et al., 1998) and bel *Drosophila* homologs of *MrDb* (Grandori et al., 1996) and *DDX3* (Liao et al., 2019) respectively. Interestingly, few of these proteins, such as pit (Zaffran et al., 1998), modulo (mod) (Perrin et al., 2003), Nnp (Nnp1) (Hulf et al., 2005), and peter pan (ppan) (Zielke et al., 2022), have been previously identified as putative direct targets of MYC specifically in the context of controlling cell growth and proliferation.

This analysis also found a highly represented cluster containing Ythdc1 (YTH domain RNA Binding Protein C1), Flacc (Fl(2)d-associated protein), and splenito (nito). Ythdc1 is a conserved nuclear m^6^A ‘reader’ protein that mediates the incorporation of methylated mRNAs into the nuclear export pathway (Roundtree et al., 2017;Shi et al., 2021). Interestingly, Flacc was found to be associated with female lethal (Fl(2)d), a protein homolog of Wilms’-tumor-1-associated protein (WTAP) (Penn et al., 2008), that was isolated in complexes with Snf (Penn et al., 2008), a component of U1 and U2 small nuclear ribonucleoproteins (snRNPs) that contained U2AF50, U2AF38, and U1-70K (small nuclear ribonucleoprotein 70K), which function in the regulation of the spliceosome. Notably, we observed an enrichment of the U2A proteins in our analysis (Supplementary, Table1), suggesting that NOC1 may play a key role in RNA splicing by linking the U1 snRNP particle to regulatory RNA-binding proteins and in the control of nuclear export via Ythdc1.

### NOC1 expression in the nucleolus increases upon MYC induction

We previously showed that endogenous NOC1 colocalizes with fibrillarin in the nucleolus (Destefanis et al., 2022). Here, we confirm the co-localization of endogenous NOC1-GFP, expressed as GFP fusion protein (NOC1-GFP) under its endogenous promoter (Kudron et al., 2018) with fibrillarin. This is clearly seen in the gigantic nucleolus of the salivary gland cells (Figure 3A-C) and also in the nucleolus of cells from the wing imaginal disc (Figure 3D-F). Furthermore, expression of MYC in cells of the wing imaginal disc, using *rotund-Gal4* promoter, increases the level of NOC1-GFP in the nucleolus and also its localization with fibrillarin (Figure 3G-I), which is a direct transcriptional target of MYC.

**Figure 3.**
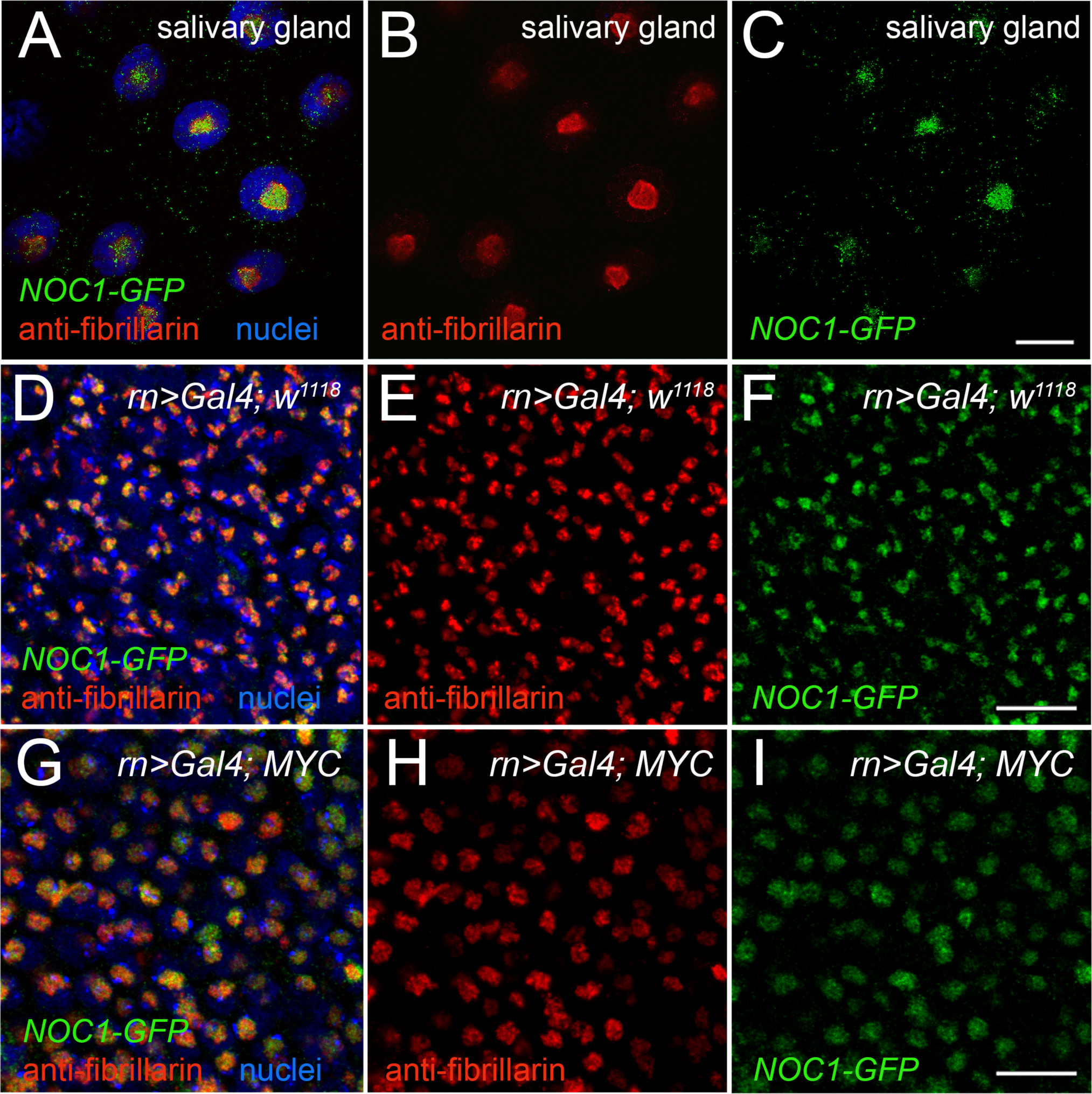
NOC1 nucleolar expression increases with MYC induction. (A-C) Confocal images of the cells from the salivary glands showing the endogenous expression of fibrillarin, visualized by immunofluorescence with anti-fib antibody (B) and NOC1-GFP nucleolar expression (C). In (A) is shown their colocalization, with nuclei stained in blue. (D-F) Cells of the wing imaginal discs show endogenous fibrillarin (E) and NOC1-GFP expression (F). Their colocalization is shown in D. (G-I) Cells of the wing imaginal discs expressing *UAS-MYC,* using the *rotund-Gal4* promoter, are stained for fibrillarin (H) and NOC1-GFP is visualized by GFP (I). In (G) is shown their merged images with nuclei stained with Hoechst (blue). Note that the nucleolus size increases by MYC expression (Grewal et al., 2005), compare G with D, which is also visible in Figure 6A. Scale bars in figures A-C represent 20 μm, and in figures D-I 10 μm.

### NOC1 overexpression induces the formation of large nuclear granules and enlarged nucleoli that co-localize with fibrillarin

We previously reported that ectopic expression of NOC1 results in nucleolar morphology changes (Destefanis et al., 2022). To analyze how the ectopic expression of NOC1 could influence nucleolar morphology, we overexpressed the HA-tagged version of NOC1 in cells of the wing imaginal discs using the *engrailed-Gal4* promoter. *Engrailed* is expressed in both the columnar epithelium forming the wing imaginal disc and in the giant cells of the peripodium, a squamous epithelium adjacent to the columnar epithelium of the wing discs (Pallavi and Shashidhara, 2005;Smith-Bolton, 2016). Analysis of NOC1 expression in these cells, by immunostaining using an anti-HA antibody, revealed in the nucleus the presence of large granules containing HA-NOC1 and an enlargement of the size of the nucleolus, where NOC1 is visibly expressed. The granules are more easily distinct and visible in the peripodium because of the gigantic size of these cells (Figure 4A and B) and with a lower resolution also in cells of the wing imaginal discs (Figures 4G and H). HA-NOC1 expression colocalizes with fibrillarin mainly in the nucleolus (Figure 4A and G), while in the granules, its expression was very low but detectable, particularly in the cells of the peripodium (Figure 4A). Co-expression of NOC1 with NOC1-RNAi visibly reduced both HA-NOC1 and the formation of the abnormal enlarged structures expression in both types of cells (Figure 3E and 3K). At the same time, the levels of fibrillarin in the nucleolus did not significantly change upon expression of NOC1-RNAi (compare Figure 4C with 4F and 4I with 4L).

**Figure 4.**
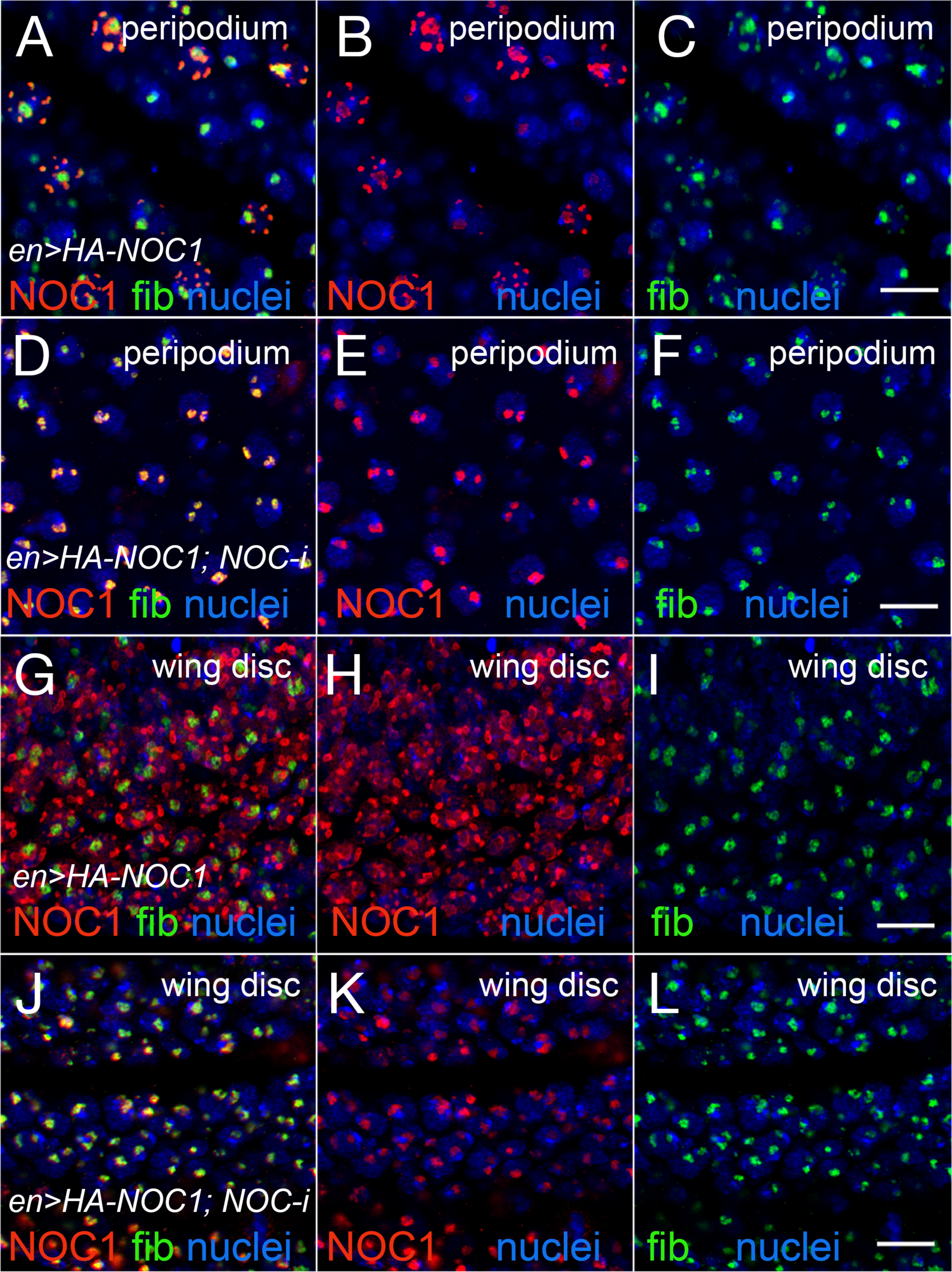
Expression of NOC1 induces extra nucleolar granules and enlargement of the nucleolus. (A-C) Confocal images of cells of the peripodium expressing HA-NOC1 alone or with NOC1-RNAi (D-F) using the *engrailed* promoter. (G-L) Images of cells from the imaginal disc expressing NOC1 alone (G-I) or with NOC1-RNAi (J-L). NOC1 and fibrillarin expression are visualized by immunofluorescence using anti-HA (red) and anti-fibrillarin (green) antibodies, respectively. Hoechst is used to visualize the nuclei. Scale bars represent 10 μm.

### NOC1 colocalizes in the nucleolus with Nucleostemin1 (Ns1) and is required for nucleolar localization of Ns1

In the analysis of proteins that can functionally interact with NOC1, we identified Nucleostemin 1 (Ns1) (Lo and Lu, 2010), a nucleolar protein necessary for the transport of the 60S subunit that shuttles between the nucleolus and the nucleoplasm, and essential for the nucleolar organization (Rosby et al., 2009). To investigate whether NOC1 interacts with Ns1, we first analyzed their co-localization. Ns1-GFP (*UAS-Ns1-GFP*) was ectopically expressed using the *patched-Gal4* promoter (Vegh and Basler, 2003) alone or with *HA-NOC1*. These data showed that when Ns1-GFP is expressed alone, it is primarily nucleolar (Figure 5AB and D) and colocalized with HA-NOC1, both in the large granules and in the enlarged nucleoli (Figure 5B), suggesting that both proteins may work together as part of a larger complex or pathway. Furthermore, we co-expressed Ns1-GFP with NOC1-RNAi, and we observed significant alterations in the subcellular localization of Ns1-GFP that, in the majority of the cells analyzed, shifted from nucleolar (Figure 5E) to the nucleoplasm (Figure 5G). These data suggest that NOC1 is required for the correct nucleolar localization of Ns1, implying that NOC1 may play a role in guiding or stabilizing Ns1 within the nucleolus (MODEL).

**Figure 5:**
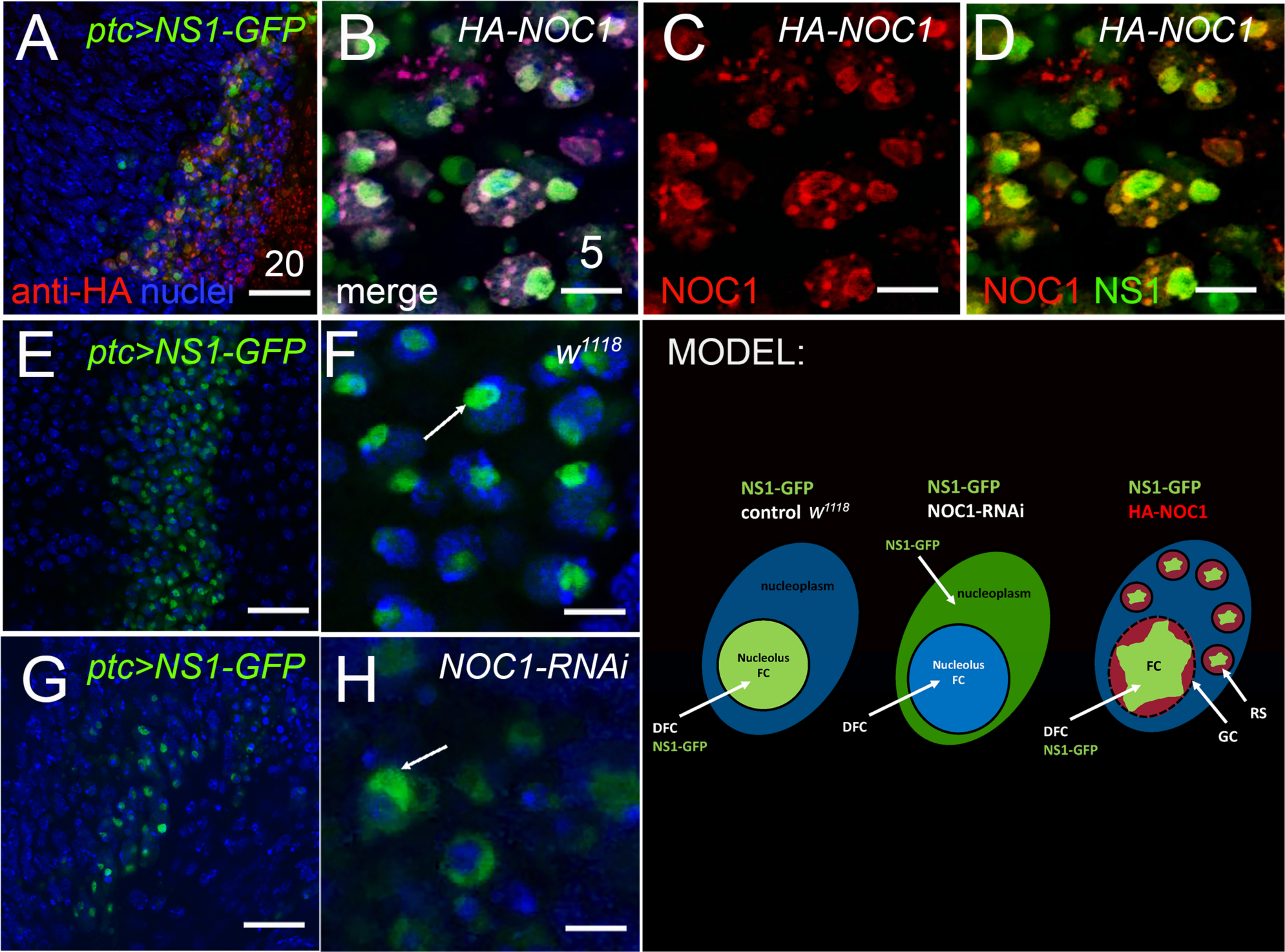
NOC1 colocalization with Nulceostemin1 (Ns1) is necessary for its nucleolar localization. Confocal images of cells from the wing imaginal disc expressing NOC1 (*UAS-HA-NOC1*) and Ns1 (*UAS-Ns1-GFP*) using the *patched-Gal4* promoter. (A-D) (A) shows a low resolution of the imaginal disc where patched is expressed as a stripe of cells along the Dorsal Ventral axis (JohnsonGrenier and Scott, 1995), as visible by GFP expression. HA-NOC1 localization is visualized by immunofluorescence using anti-HA antibodies, while Ns1 was detected by the GFP expression of the fusion protein; nuclei are stained in blue by Hoechst. The co-localization is shown in (B) as merged while in (C) the HA-NOC1 expression and in (D) Ns1-GFP and HA-NOC1 co-localization. (E) Confocal image of *ptc>Ns1-GFP* expression in the wing imaginal disc. (F) higher magnification of staining in E showing Ns1 localization in the nucleolus (Arrow). (G) Confocal image of *ptc>Ns1-GFP; UAS-NOC1-RNAi* expression in the wing imaginal disc. (H) higher magnification of staining in G showing Ns1 localization mainly in the nucleoplasm (Arrow). Nuclei are stained in blue. Scale bars in A-E and G represent 20 μm, and B-C-D-F and H represent 5 μm. MODEL suggesting the functional interaction of Ns1 with NOC1 in the nucleolus and describing the nucleolus organization as FC: Fibrillar Center, DFC: Dense Fibrillar Components, GC: Granular Center (LamTrinkle-Mulcahy and Lamond, 2005). GS: Granular Structures visualized by NOC1 overexpression.

### Coexpression of NOC1 with MYC increases the formation of abnormal structures and granules in the nucleus

We then analyzed if increasing the rate of protein synthesis by overexpressing MYC could somehow reduce NOC1 granules, assuming that they might function as storage of ribosomal factors produced in excess by NOC1 overexpression. Contrary to our hypothesis, we found that co-expression of MYC, which alone increased the nucleolar size, visualized with anti-fibrillarin antibodies (Figure 6A), and enhanced further the size of the aberrant structures containing HA-NOC1 (Figure 6B), increasing the nucleolar shape (Figure 6C), suggesting that MYC enhances NOC1 activity in the nucleolus, highlighting the complex interplay between these two factors and their effects on nucleolar homeostasis.

**Figure 6:**
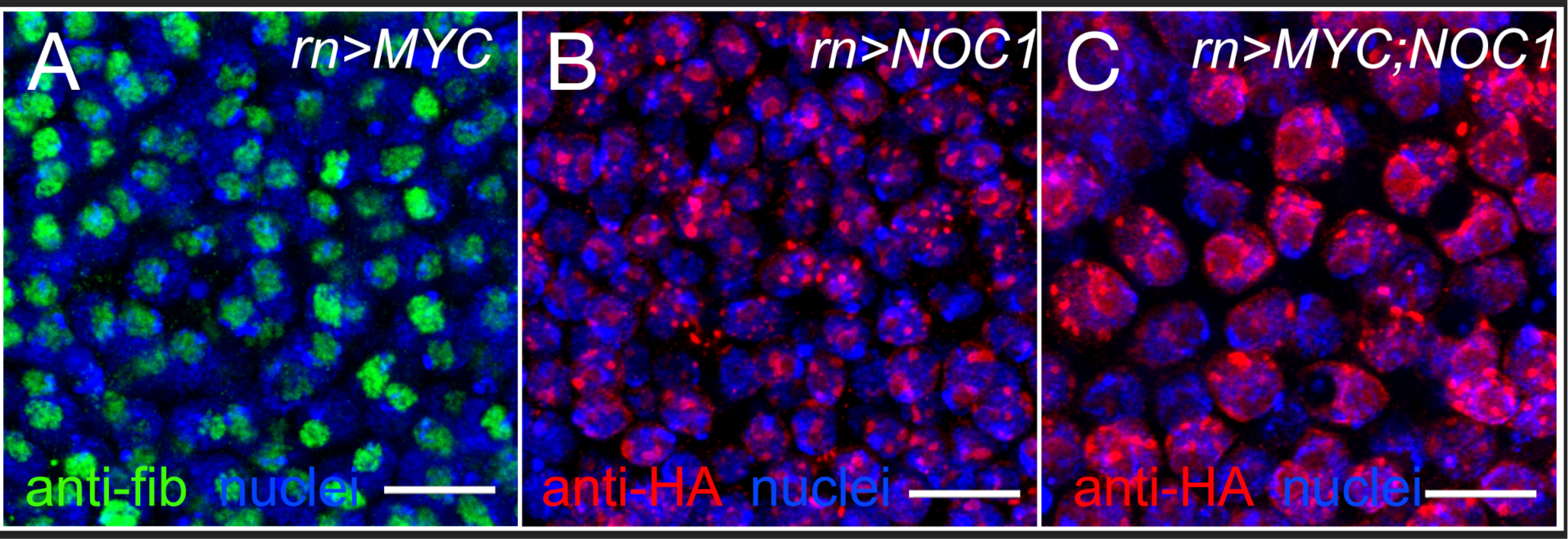
Expression of NOC1 and MYC enhances nucleolar morphology. Confocal images of cells from the wing imaginal disc expressing (A) MYC (*UAS-MYC*) (B) NOC1 (UAS-HA-NOC1), or both (C) using the *rotund-Gal4* promoter. Fibrillarin is visualized in A by immunofluorescence using anti-fibrillarin antibody (green); (B-C) immunofluorescence of NOC1 expression using anti-HA antibodies (red), nuclei are stained using Hoechst and visualized in blue. Scale bars represent 10 μm.

## Discussion

The nucleolus is a critical subcellular compartment involved in ribosome biogenesis, and proteins like NOC1 play essential roles in this process. The conservation of NOC1 function across these diverse organisms, from yeast (*S.ce*) *Arabidopsis, Drosophila* (Milkereit et al., 2001;Li et al., 2009;de Bossoreille et al., 2018;Destefanis et al., 2022) and to some extent humans (Barbieri et al., 2017)(our unpublished data), indicates the fundamental role of this nucleolar factor in controlling basic and essential processes during ribosome biogenesis.

We have recently characterized the function of the sole nucleolar *NOC1* gene in *Drosophila* and show that it is necessary for proper rRNA processing and maturation, while its downregulation reduces protein synthesis and is detrimental to organ and animal growth (Destefanis et al., 2022). Here, we characterized NOC1 as a *bona fide* MYC target gene and demonstrated that NOC1 is transcriptionally induced through a functional MYC-binding E-box sequence in the NOC1 promoter region (Figure 1). We then analyzed NOC1 interactome by MS analysis (Supplementary list1) to identify how NOC1 functions in controlling ribosomes and in relation to MYC. These data reveal that NOC1 is in a complex with the nucleolar proteins NOC2 and NOC3, confirming previous data in yeast, and probably forms functional heterodimers necessary for the transport of the large ribosomal subunit during ribosome maturation (Milkereit et al., 2001;Hierlmeier et al., 2013). Our data also evidence an enrichment in NOC1-IPs of other nucleolar proteins, many of them such as fib, mod, nnp1, have been previously characterized as direct MYC’s targets (Perrin et al., 2003;Hulf et al., 2005). In support of this last observation, we also found that in response to MYC, NOC1 expression and localization within the nucleolus is significantly increased, suggesting a direct functional response between MYC and NOC1 activities in this organelle. Notably, NOC1 overexpression leads to the formation of large nuclear granules and enlarged nucleoli, which co-localizes with nucleolar fibrillarin and Ns1. Additionally, we demonstrate that NOC1 expression is necessary for Ns1 nucleolar localization, suggesting a role for NOC1 in maintaining nucleolar structure. Finally, the co-expression of NOC1 and MYC enhances the formation of abnormal structures within the nucleus containing NOC1, outlining another aspect where NOC1 and MYC activities may cooperate or be additive in controlling nucleolar dynamics.

Furthermore, our study also highlights NOC1 interaction with proteins relevant for RNA processing, modification, and splicing. Indeed, we found highly represented Ythdc1 and Flacc (Fl(2)d-associated protein) and spenito (nito), the flies homolog of the nucleolar large ribosomal subunit (60S) assembly factor RBM28 (Bryant et al., 2021). Notably, all these proteins are part of the mechanism that mediates N6-methyladenosine (m6A) methylation of mRNAs (Shi et al., 2021;Deng et al., 2023). Ythdc1 is a conserved nuclear m^6^A ‘reader’ protein that mediates the incorporation of methylated mRNAs for their nuclear export (Roundtree et al., 2017;Shi et al., 2021). Flacc is a component of the complex that mediates N6-methyladenosine methylation of mRNAs essential for mRNA splicing efficiency of pre-mRNA targets and a key regulator of Sxl (Sex-lethal) pre-mRNA splicing (Knuckles et al., 2018). Flacc is in complex with female lethal (Fl(2)d), the *Drosophila* homolog of Wilms’-tumor-1-associated protein (WTAP) a component of human spliceosome (Zhou et al., 2002), and with Snf a component of U1/U2 small nuclear ribonucleoproteins (snRNPs) that contained U2AF50, U2AF38, and U1-70K necessary for splicing reaction of pre-mRNAs (Penn et al., 2008). Interestingly, we found an enrichment of these proteins in our analysis (Supplementary Table 1). Additionally, our data may suggest a potential link between NOC1 and snRNPs involved in regulating RNA-binding proteins and controlling mRNA nuclear export via m6A-dependent modifications by Ythdc1. This part highlights the complex and interconnected processes involved in gene expression regulation, from mRNA splicing to modifications. However, how NOC1 may control or be part of these mechanisms is still unclear.

Our previous analysis directly assessed the impact of NOC1 on pre-rRNA processing and cleavage and showed that its reduction induced an accumulation of pre-rRNA precursors (ITS1 and ITS2) (Destefanis et al., 2022). Similar data were found for the NOC1 homolog in yeast (Noc1p) using genetic screens and proteomic studies (Hierlmeier et al., 2013;Lebaron et al., 2013;Khoshnevis et al., 2019). However, we should comment on some crucial differences in the protein-interactome from our experiments and those in yeast. Few reports in yeast annotated the Noc1p protein associated with Rrp5 (Ribosomal RNA Processing 5), a factor crucial for ribosome assembly that mediates the cleavage of the 35S pre-rRNA into the 18S rRNA, which is a critical step in the production of the small ribosomal subunit (Hierlmeier et al., 2013;Lebaron et al., 2013) and with Rcl1 (Ribosomal RNA Cleavage 1), another enzyme with a role in rRNA cleavage and processing (Khoshnevis et al., 2019). Both these proteins are conserved in flies. However, we did not find them in our NOC1-interactome analysis, even though Rrp5 was found in yeast bound to the pre-rRNA region of the ITS1 (Internal Transcriber Spacers-1) using protein crosslinking following by RNase treatment (Lebaron et al., 2013), and to interact with Noc1p and Noc2p (Hierlmeier et al., 2013) with a Noc1p-TAP purification system. We can explain these differences by hypothesizing that either the levels of Rrp5 and Rcl1 expressions are low in larvae compared to yeast or the use of different techniques and timing of purification of the protein used as bait in yeast compared to ours, i.e., during specific phases of RNA maturation and using Noc1p-TAP purification systems (Sailer et al., 2022). However, in the NOC1-interactome, we found NOC2 and NOC3, along with Nop53 (Rrp9) among others, described part of the Noc1p-yeast complex (Ohmayer et al., 2013), highlighting the significance of our preliminary studies in flies. It’s important to acknowledge that studying the precise protein-interactome of NOC1 in in vivo can be challenging, and experimental conditions can limit the interpretation of results. In our case, conducting experiments at a single time point and under standard immunoprecipitation conditions may provide valuable insights into protein interactions but might not fully capture the dynamic and context-dependent nature of different NOC1’s functions.

We found that NOC1 overexpression forms large granular structures containing NOC1, along with fibrillarin and Nuleostemin1 (Figures 4 and 5). At the moment, we do not know the nature of these granules. We could hypothesize that these HA-NOC1 granules work as dynamic and multifunctional structures regulating RNA metabolism and gene expression, including rRNA processing and transcription. These may include RNA stress granules formed during stress conditions to protect mRNAs from degradation or to control their translation (PutnamThomas and Seydoux, 2023). This hypothesis is supported by our data that identify proteins of the DEAD-box RNA helicases family, such as pea/DXH8 and CG8611 pit, bel kurz, previously identified as components of RNA stress granules (Campos-Melo et al., 2021). This idea may also support the mechanism by which the abnormally large structures containing NOC1 and induced when MYC is overexpressed are the result of their synergistic effect in promoting cellular stress induced by a high protein synthesis or dysfunctions caused by the combination of MYC and NOC1 targets. Overexpression of MYC can lead to increased demand for ribosome biogenesis, and the presence of abnormal ribosomal intermediates due to NOC1 dysregulation can exacerbate this stress. This can result in nucleolar stress, activation of cellular stress responses, and potentially contribute to the insurgence of diseases.

Abnormal structures or extra nucleoli have significant implications in human diseases, particularly in cancer, where dysregulation of nucleolar functions is a hallmark of the disease (Orsolic et al., 2016;Penzo et al., 2019), and in ribosomapathies a class of rare genetic diseases characterized by mutations in ribosomal proteins or components that impaired RNA translation associated with various clinical manifestations, including bone marrow failure, developmental disorders and an increased risk of cancer (Farley-BarnesOgawa and Baserga, 2019;Kampen et al., 2020).

Finally, a few words about the human homolog of NOC1, called CEBPz (CCAAT/enhancer-binding protein zeta), a transcription factor so far associated with certain types of tumors. Notably, in acute myeloid leukemia (AML), CEBPz was shown to promote the m^6^A modification of target mRNA transcripts, enhancing their translation (Barbieri et al., 2017;HongXu and Lee, 2022). Thus, overexpression or downregulation of CEBPz in humans may also affect RNA processing, leading to defective translation. In support of this idea, the human gene rbm28, which we found in the NOC1 interactome, is responsible for the ribosomapathy-ane syndrome (Bryant et al., 2021), a rare genetic disorder caused by aberrant splicing in *RBM28* pre-mRNA. This, together with other indirect information on the potential role of NOC1/CEBPz in controlling alternative splicing, highlights the potential role of the human counterpart in the control of nucleolar processes that may cause genetic disorders.

Our research uses *Drosophila*, a simple and accessible model system, to identify novel conserved mechanisms to better understand MYC activity and its targets, including NOC1, in the context of RNA translation and ribosome biogenesis. The ultimate goal would be to identify specific targets within the translation machinery that small molecules or drugs can modulate for use in disease therapies.

## Supporting information

List Cluster 1

## Notes

### Competing Interest Statement

The authors have declared no competing interest.

